# Selective quality control of mistargeted mitochondrial proteins at the endoplasmic reticulum

**DOI:** 10.64898/2026.08.10.743879

**Authors:** Yuichi Tsuchiya, Nikita Sergejevs, Maria Emilia Dueñas, Mike Renne, Elena Navarro-Guerrero, Matthias Trost, Pedro Carvalho

**Author notes:** co-corresponding authors: Correspondence should be sent to: Pedro Carvalho and Yuichi Tsuchiya.

## Abstract

In eukaryotic cells, the function of each organelle depends on its unique protein composition. Protein targeting errors threaten organelle identity and function, yet how mistargeting errors are detected and resolved remains poorly understood. Here, we show that mitochondrial import stress drives widespread rerouting of mitochondrial proteins to the endoplasmic reticulum (ER), with strong enrichment for hydrophobic oxidative phosphorylation (OXPHOS) components. Using proximity proteomics, a split-fluorescence reporter system, and genome-wide CRISPR screening, we find that mistargeted proteins partition into distinct classes with divergent fates, ranging from stable ER residence to rapid degradation by ER-associated degradation (ERAD). The clearance of these mislocalized proteins involves partially redundant ERAD branches, with the ubiquitin ligase MARCHF6 playing a central role. Together, these findings establish the ER as a key organelle for handling mistargeted mitochondrial proteins and reveal how ER quality control maintains proteostasis during mitochondrial dysfunction.

## Introduction

Accurate protein localization is essential for the organization and function of eukaryotic cells. This is particularly critical for membrane proteins, whose insertion into the correct lipid bilayer supports organelle identity and function. Protein targeting pathways, such as those mediated by signal recognition particle (SRP) for the endoplasmic reticulum (ER) or dedicated import machineries for mitochondria, ensure that newly synthesized proteins reach their appropriate destination (Hegde and Keenan, 2022; Endo and Wiedemann, 2025). However, targeting is inherently error prone (Guna and Hegde, 2018; McKenna and Shao, 2023). As a result, cells rely not only on dedicated targeting mechanisms but also on robust quality control systems that monitor protein localization and eliminate proteins that fail to reach their correct compartment (McKenna and Shao, 2023; Krshnan et al., 2022; Christianson and Carvalho, 2022).

Mitochondria present a major challenge for protein targeting fidelity. Although they contain a small genome, the vast majority of mitochondrial proteins are encoded in the nucleus, synthesized in the cytosol, and imported mostly post-translationally (Endo and Wiedemann, 2025; Rackham and Filipovska, 2022). This process depends on targeting signals such as N-terminal mitochondrial targeting sequence (MTS) and on the coordinated action of cytosolic chaperones (Bykov et al., 2020, 2022). Under physiological conditions, import is highly efficient, but it can be compromised by mutations, metabolic stress, or defects in the import machinery, leading to the accumulation of non-imported precursor proteins in the cytosol (Krämer et al., 2023; Nowicka et al., 2021; Pfanner et al., 2025). These precursor proteins are often aggregation-prone and potentially toxic, necessitating efficient surveillance and clearance mechanisms (Song et al., 2021; Liu et al., 2025; Wrobel et al., 2015; Boos et al., 2019).

Emerging evidence indicates that mislocalized mitochondrial proteins are not exclusively handled in the cytosol but can be rerouted to other organelles, most notably the ER (McKenna et al., 2020; Matsumoto et al., 2019; Xiao et al., 2021; Vitali et al., 2018; Gamerdinger et al., 2015; Hansen et al., 2018). In yeast, several mitochondrial membrane proteins have been shown to localize to the ER when import is impaired (Xiao et al., 2021; Hansen et al., 2018), in some cases via pathways normally dedicated to ER targeting (McKenna et al., 2022; Vitali et al., 2018). In mammalian cells, however, the extent to which mitochondrial proteins are mistargeted to the ER and the general principles governing this process remain poorly defined.

Protein targeting to the ER is itself a highly regulated and competitive process. While SRP recognizes signal sequences and directs nascent chains to the ER co-translationally, ribosome-associated factors such as the nascent polypeptide-associated complex (NAC) prevent inappropriate engagement of mitochondrial and other non-secretory proteins, thereby modulating targeting fidelity (Jomaa et al., 2022; Muthukumar et al., 2024; Lee et al., 2026). Perturbations in this balance, such as those arising from the accumulation of non-imported mitochondrial precursors, could increase the likelihood that these proteins are aberrantly captured by ER targeting pathways, particularly if they contain hydrophobic segments (McKenna and Shao, 2023; Gamerdinger et al., 2015; Pfanner et al., 2025).

Once proteins reach the ER, their fate is determined by ER quality control systems such as ER-associated degradation (ERAD). This process identifies aberrant or mislocalized proteins, ubiquitinating them, and targeting them for proteasomal degradation (Christianson et al., 2023; Christianson and Carvalho, 2022). Multiple ERAD branches, including those mediated by MARCHF6, HRD1 and RNF145, differ in substrate specificity and recognition mechanisms (Krshnan et al., 2022; Sergejevs and Carvalho, 2025). However, how these pathways handle proteins mistargeted from other organelles, and whether such substrates are processed similarly remains largely unknown.

An important unresolved question is whether all mistargeted proteins are treated equivalently by the ER quality control machinery. Some studies suggest that certain mitochondrial proteins can adopt alternative localizations or remain stable outside mitochondria (Yogev and Pines, 2011; Friedman et al., 2018) raising the possibility that the fate of mistargeted proteins is heterogeneous and dictated by intrinsic features such as topology, hydrophobicity, or sequence context. Defining the spectrum of mistargeted proteins and understanding how their properties influence their recognition and processing is therefore critical for elucidating how cells maintain proteome integrity.

Here, we set out to systematically characterize the mistargeting of mitochondrial proteins to the ER and to define the mechanisms that govern their fate. To this end, we combined ER-targeted proximity labeling with quantitative proteomics to identify mitochondrial proteins that localize to the ER under import stress. We further developed a split-fluorescence reporter system to monitor mistargeting and degradation at the level of individual substrates and performed genome-wide CRISPR screens to define the cellular machinery involved. Using this approach, we show that mitochondrial import stress induces widespread and selective rerouting of mitochondrial proteins to the ER, where they are partitioned into distinct classes and cleared by a network of ERAD pathways centered on the ubiquitin ligase MARCHF6.

## Results

### Global protein mistargeting to the ER during mitochondrial import stress

Certain mitochondrial proteins were shown to associate with the ER when their import into mitochondria is compromised (Gamerdinger et al., 2015; Shakya et al., 2021; Hansen et al., 2018). However, the extent of this ER relocalization and how general is this phenomenon across the mitochondrial proteome remains ill defined.

To study protein mistargeting to the ER in an unbiased manner, we used a proximity biotinylation approach. To this end, we targeted the ascorbate peroxidase APEX2 with a C-terminal ER retention KDEL peptide (Hung et al., 2016) to the ER lumen (ER-APEX2-K) by fusing it to the N-terminal signal sequence of human BiP. We hypothesized that, under conditions of mitochondrial import stress, mitochondrial proteins targeted to the ER would become substrates of ER-APEX2-K and biotinylated (Figure 1A). Moreover, we expected ER-APEX2-K to primarily modify mitochondrial precursor proteins, containing their targeting signal and whose removal occurs only in the mitochondrial matrix upon their import (Figure 1A). To trigger mitochondrial import stress, we used two agents, alone or in combination: the ionophore valinomycin, which blocks membrane-potential-dependent precursor import, and ISRIB, which inhibits the integrated stress response, thereby preventing stress-induced attenuation of cytosolic translation and sustaining the supply of newly synthesised precursors. In each case the proteasome inhibitor bortezomib was included to stabilise mistargeted precursors and enable their detection. As a proof of principle for this assay, we analysed endogenous COX4I1, a nuclear encoded assembly factor of complex IV of oxidative phosphorylation (OXPHOS) both in the absence and presence of mitochondrial import stress (Vercellino and Sazanov, 2022). In unperturbed cells, the COX4I1 precursor was undetectable, and the mature protein was not modified by ER-APEX2-K, as expected (Figure 1B). In contrast, under conditions of mitochondrial import stress in combination to proteasome inhibition, led to the accumulation of COX4I1 precursor. Under these stress conditions the precursor, but not mature COX4I1, was mistargeted to the ER and modified by ER-APEX2-K (Figure 1B). Similar conditions also resulted in ER mistargeting of other mitochondrial precursor proteins such as DHFR and OXA1L (Figure 1C). Importantly, abundant cytosolic proteins, such as Tubulin, were not modified by ER-APEX2-K indicating the specificity of labelling. In contrast, ER proteins, like Calnexin, were modified irrespective of the presence of mitochondrial stress, as expected (Figure 1C). These data indicate that mitochondrial import stress leads to mistargeting of mitochondrial precursors to the ER and demonstrate that proximity biotinylation by ER-APEX2-K is a robust tool to study this process.

**Figure 1.**
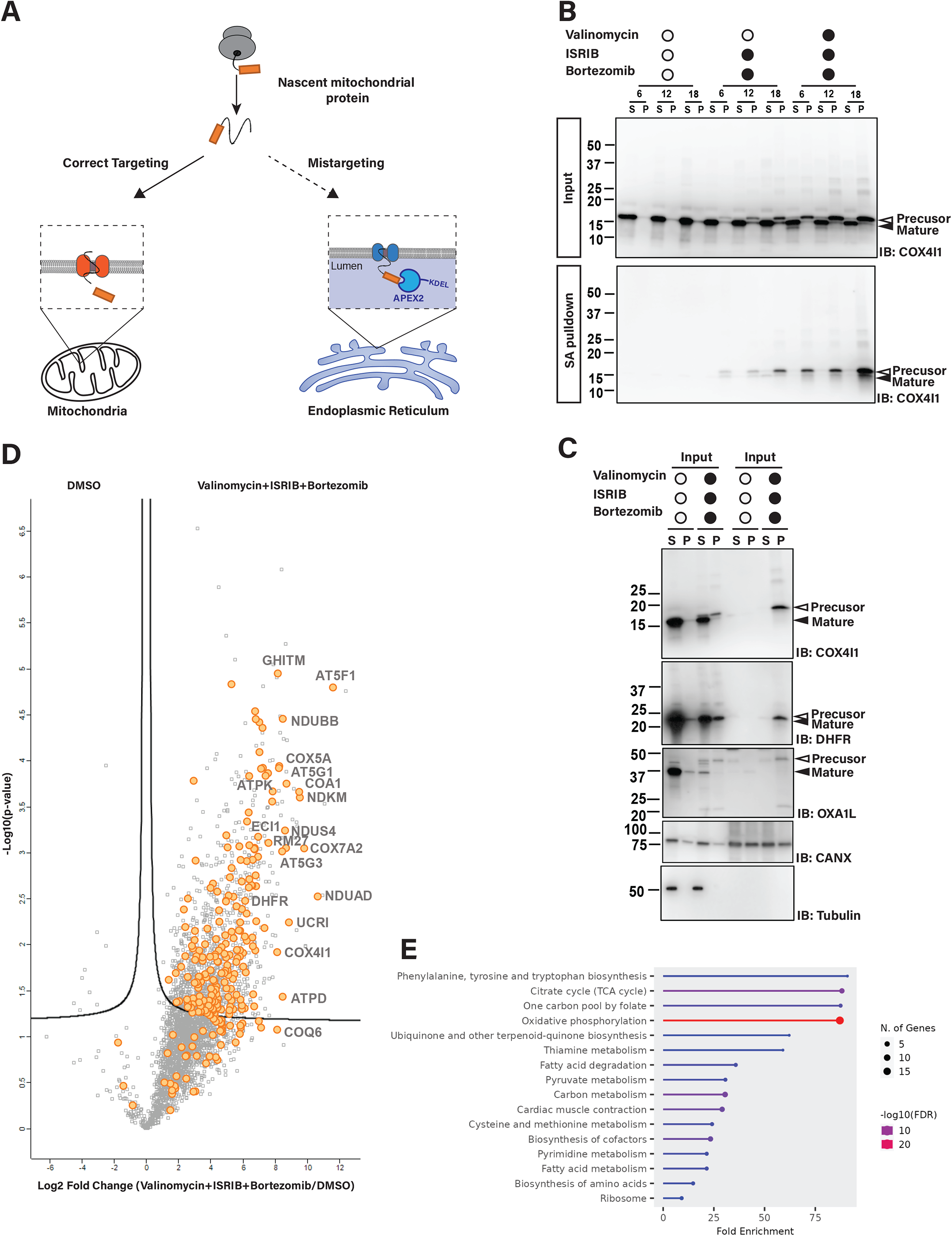
Global protein mistargeting to the ER during mitochondrial import stress. **(A)** Schematic representation of APEX2-based proximity labeling assay for unbiased detection of ER-mistargeting upon mitochondrial import stresses. **(B)** Biotinylation of endogenous COX4I1 by ER targeted APEX2 upon mitochondrial import stress. HEK293T cells stably expressing ER-APEX2-KDEL were treated with 1 μM valinomycin, 400 nM ISRIB and/or 1 μM bortezomib for the indicated times. Cell lysates were subjected to centrifugation and supernatant (S) and pellet (P) fractions were analysed by SDS-PAGE before and after streptavidin pulldown. Note that precursor COX4I1 proteins is specifically biotinylated and precipitated by streptavidin beads. COX4I1 was detected with an anti-COX4I1 antibodies. **(C)** Selective biotinylation of mitochondrial precursor proteins upon import stress. Cells were treated and analysed as in (B). **(D)** Proteins modified by ER-APEX2-KDEL and precipitated by streptavidin beads upon mitochondrial import stress as detected by mass spectrometry. The x axis shows the log2 fold change under import stress versus unstress control cells; the y axis shows the −log10 p value estimated by the SignificanceB analysis (Cox and Mann, 2008). Mitochondrial proteins are indicated in orange. **(E)** GO term enrichment among the top 200 proteins in Fig. 1D as analysed by ShinyGo 0.85 (https://bioinformatics.sdstate.edu/go/).

To have a complete repertoire of mistargeted proteins during mitochondrial import stress, we carried out the assay described above followed by mass spectrometry to identify proteins modified by ER-APEX2-K (Figure 1D). Import stress combined with proteasomal inhibition led to a large and specific increase of mitochondrial proteins in the ER, with components of OXPHOS being particularly enriched (Figure 1E). These proteins are highly hydrophobic, many being integral to the membrane and containing multiple transmembrane segments, features that likely explain their robust ER mistargeting and modification by ER-APEX2-K. Altogether, these data indicate that mitochondrial import stress results in a widespread redirection of mitochondrial proteins to the ER, with many proteins, in particular IMM proteins such as OXPHOS components becoming exposed to the ER lumen.

### Distinct fates of mistargeted mitochondrial proteins at the ER

To study the fate of mistargeted mitochondrial to the ER, we focused on a set of OXPHOS proteins, as our mass spectrometry analysis indicated that they are particularly prone to ER mistargeting. To this end, we employed a split fluorescence reporter assay in which the N-terminal half of moxVenus fluorescent protein was targeted to the ER lumen by fusing it to the N-terminal signal sequence of human BiP and a C-terminal ER retention KDEL peptide (ER-moxV(N)-K) (Costantini et al., 2015). This construct also included mCherry fluorescent protein to facilitate detection by microscopy, western blotting and flow cytometry. In parallel, the C-terminal half of moxVenus (moxV(C)) was fused to a protein of interest, most commonly an OXPHOS component, and expressed from a doxycycline responsive locus (Figure 2A). This reporter system was highly specific, with robust complementation occurring only when both halves of moxVenus faced the ER lumen, whereas an otherwise identical acceptor exposing moxVenus(N) to the cytosol (moxV(N)-CytB5) gave little or no signal (Figure S1A). It should be noted that while the assay reports on the luminal exposure of the substrate C-terminus, it does not, by itself, define the membrane topology or insertion state of the mistargeted protein, which was not established directly.

**Figure 2.**
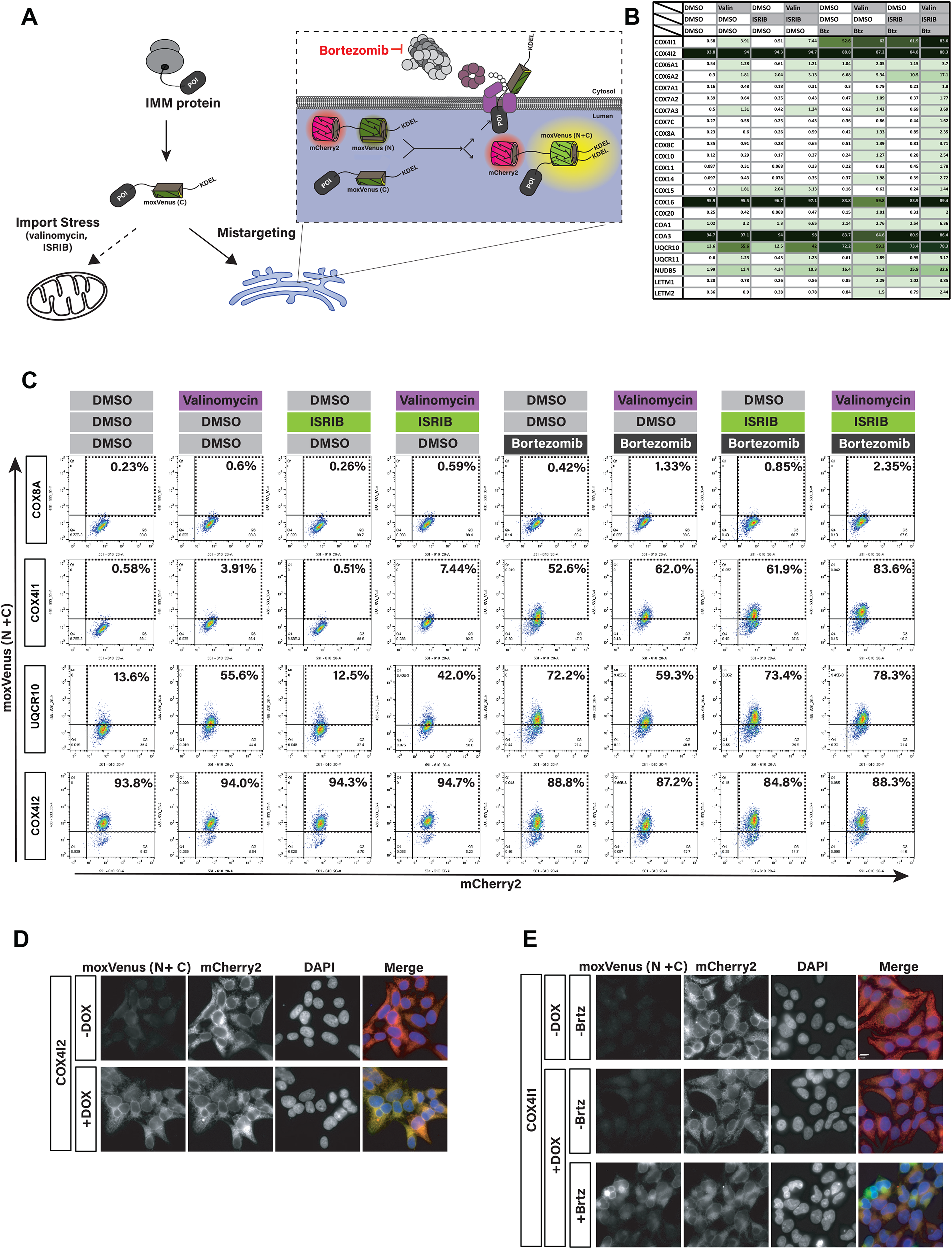
Distinct fates of mistargeted mitochondrial proteins at the ER. **(A)** Schematic representation of split-moxVenus-based strategy to detect mislocalized proteins in the ER lumen. **(B)** Percentage of cells with positive moxVenus signal as determined by flow cytometry. Rows indicate individual proteins expressed as C-terminal fusions to moxVenus(C) in cells also expressing ER-targeted mCherry-moxVenus(N)-KDEL while columns indicate the various conditions tested. **(C)** Parental cells expressing the indicated split-moxVenus reporters were subjected to the indicated treatments and analysed by flow cytometry. The percentage of moxVenus-positive cells is indicated in each case. **(D)** Immunofluorescence of HEK293T cells expressing the split moxVenus COX4I2 reporter and analysed under the indicated condition. mCherry channel shows the signal of ER targeted mCherry-moxVenus(N)-KDEL. **(E)** As in (D) but showing cells expressing split moxVenus COX4I1 reporter.

Using this assay, we analysed several OXPHOS components which showed distinct behaviours in terms of mistargeting and fate. Some proteins, like COX4I2, COX16 and COA3, showed robust localization to the ER lumen even in the absence of any mitochondrial import stress and their levels were unaffected by proteasome inhibition suggesting that they were stable upon ER mistargeting (Figure 2B-C).

Other proteins, such as COX4I1 and UQCR10, were barely detected in the ER under basal conditions but their ER localization became apparent upon proteasome inhibition with Bortezomib suggesting that they were subject to proteasomal degradation likely via ERAD (Figure 2 B-C). Importantly, their ER localization increased during mitochondrial import stress (Figure 2B-C) suggesting that mistargeting occurred at basal conditions and was further increased under mitochondrial stress.

To confirm the localization of COX4I1 and COX4I2 we analysed the split moxVenus complementation signal by fluorescence microscopy before and after doxycycline-induced expression. A strong moxVenus signal overlapping with ER-targeted mCherry was readily detected in COX4I2 (Figure 2D). Similarly, COX4I1 was also detected at the ER but only upon proteasome inhibition with bortezomib treatment (Figure 2E). Consistent with the FACS analysis, these data indicate that while both COX4I1 and COX4I2 can mistarget to the ER, only COX4I1 is unstable in this organelle.

### MARCHF6 has a key role in the degradation of ER mistargeted OXPHOS proteins

To identify factors that promote mistargeting to and/or clearance of OXPHOS proteins from the ER, we performed a genome wide CRISPR screen. Cells with constitutive expression of (ER-moxV(N)-K) and doxycycline-inducible expression of COX4I1-moxV(C)-KDEL (K) were transduced with the Toronto KnockOut CRISPR-Cas9 Library version 3 (TKOv3), which contains a total of 70,948 sgRNAs targeting 18,053 human genes (Hart et al., 2017). Mutant cells displaying high moxVenus fluorescence, indicative of mistargeted COX4I1, were isolated by flow cytometry at two different timepoints. Next-generation sequencing was used to sequence and quantify sgRNAs in reference and sorted cell populations (Figure 3A) and the MAGeCK algorithm (Li et al., 2014) was used to rank genes based on their effect on COX4I1 mistargeting (Figure 3B and S1B). Analysis of two timepoints allowed us to capture the contributions of both non-essential genes, which persist, and essential genes, which are initially present but depleted at later stages (van de Weijer et al., 2020).

**Figure 3.**
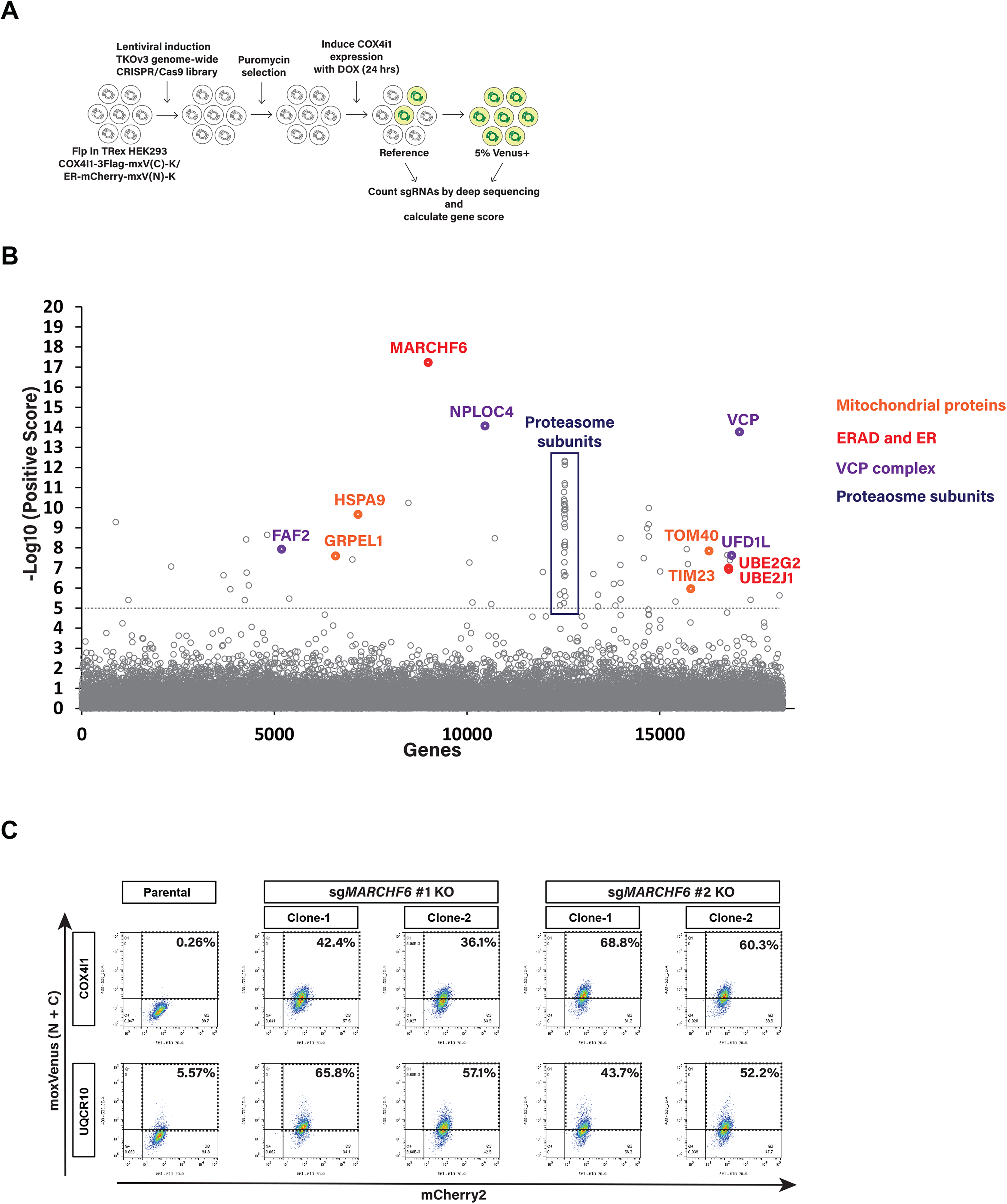
A Genome-wide CRISPR-Cas9 Screen Identifies Components Required for the degradation of mistargeted COX4I1. **(A)** Workflow of the CRISPR-Cas9 genome-wide screen. **(B)** Significance score of the genes analyzed in the screen calculated by the MAGeCK algorithm. The x axis represents the genes in alphabetical order. The y axis shows the −log(αRRA) significance value. The −log(αRRA) cutoff was arbitrarily set at 9 (dashed line). Significantly enriched genes are annotated. COX4I1 reporter expressing cells were transduced with the TKOv3 sgRNA library and the cell population with high moxVenus fluorescence levels was isolated by flow cytometry 8 days post transduction. **(C)** Depletion of MARCHF6 with 2 independent sgRNAs result in the stabilization of mistargeted mitochondrial proteins COX4I1 and UQCR10, as detected by flow cytometry of cells expressing the split-moxVenus reporter system.

Key components of the mitochondrial import machinery such as *TOM40* and *TIM23*, were among the hits, which was expected given the known effect of their depletion in inducing mitochondrial import stress (den Brave et al., 2024). Depletion of mitochondrial chaperones, such as *HSPA9*, a mitochondrial HSP70, and its nucleotide exchange factor *GRPEL1* (Morizono et al., 2024) also led to increased COX4I1 mistargeting, likely due lower protein translocation efficiency (Michaelis et al., 2022) (Figure 3B).

Strikingly, the most numerous and prominent hits were protein quality control factors, including components of the proteasome, the p97 ATPase complex, and ERAD machinery. Among these, the highest ranked was the ER-resident ubiquitin ligase *MARCHF6* as well as the conjugating enzymes *UBE2J2* and *UBE2G2*, which enable MARCHF6-dependent ubiquitination (Stefanovic-Barrett et al., 2018), suggesting a key role for this ERAD branch in the elimination of mistargeted COX4I1 (Figure 3B). Independent depletion of *MARCHF6* with additional sgRNAs led to the accumulation of mistargeted COX4I1 and UQCR10, another mitochondrial protein suggesting this ubiquitin ligase plays a broader role in the ERAD of mistargeted OXPHOS proteins (Figure 3C).

To investigate the role of MARCHF6 in the degradation of mistargeted mitochondrial proteins, COX4I1 was used as a model substrate. Western blotting analysis showed that ablation of *MARCHF6* resulted specifically in the accumulation of COX4I1 precursor protein (Figure 4A, S2A). Re-expression in *MARCHF6* KO cells of wild type *MARCHF6* but not the *MARCHF6* C9A mutant, which lacks ubiquitin ligase activity, reversed this phenotype indicating that MARCHF6 ubiquitin ligase activity is critical for the clearance of mislocalized COX4I1(Figure 4B-D). The effect was specific as *MARCHF6* deletion or overexpression had no effect on the levels of COX4I2, a protein that remains stable when mislocalized to the ER (Figure 4C and D). Consistent with these results, MARCHF6 specifically interacted with the COX4I1 precursor but not with its mature form. This interaction was even more prominent with MARCHF6 C9A mutant, which binds substrates efficiently but is defective in their ubiquitination (Figure 4E).

**Figure 4.**
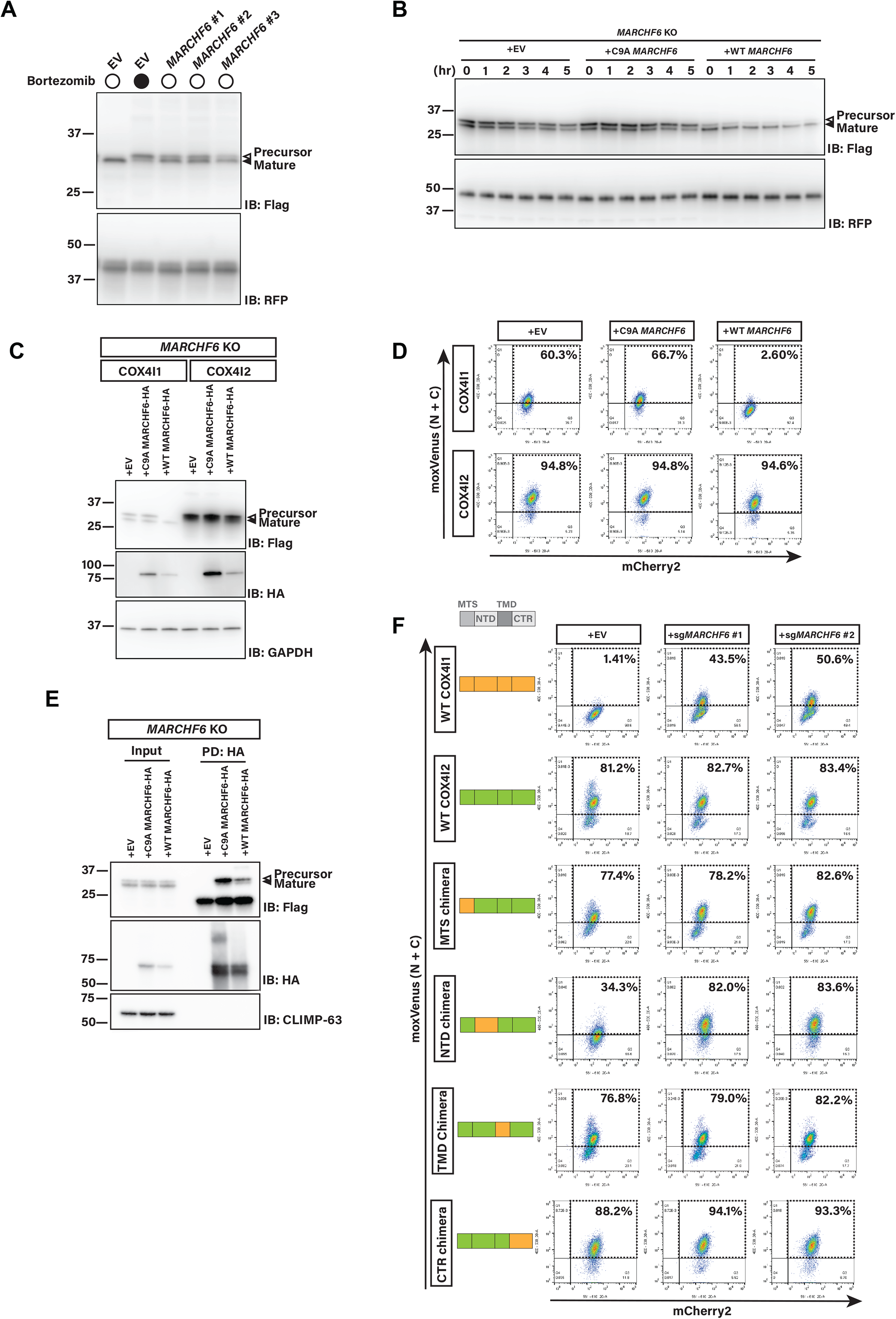
MARCHF6 has a key role in the degradation of ER mistargeted COX4I1. **(A)** Depletion of MARCHF6 or proteasome inhibition result in specific accumulation of COX4I1 precursor. HEK293 cells with the split-moxVenus system expressing COX4I1-FLAG-moxVenus(C) and ER targeted mCherry-moxVenus(N)-KDEL were subjected to to the indicated treatments and extracts were analysed by SDS-PAGE and immunoblotting. COX4I1-FLAG-moxVenus(C) was detected with anti-FLAG antibodies. mCherry was used as loading control and detected with anti-RFP antibody. **(B)** MARCHF6 promotes the degradation of COX4I1 precursor as detected in a cycloheximide chase experiment. Extracts MARCHF6 KO cells expressing the indicated variants were collected at the indicated timepoints upon inhibition of protein synthesis with cycloheximide and analyzed by SDS-PAGE and immunoblotting. COX4I1-FLAG-moxVenus(C) was detected with anti-FLAG antibodies. mCherry was used as loading control and detected with anti-RFP antibody. **(C)** MARCHF6 ubiquitin ligase activity affects COX4I1 precursor while it has no impact on COX4I2 precursor protein. HEK293T cells lacking MARCHF6 and expressing HA tagged of the indicated MARCHF6 variant as well as the COX4I1 or COX4I2 moxVenus-split system were analyzed by western blot. COX4I proteins were detected with anti-FLAG antibodies. MARCHF6 variants were detected with anti-HA antibodies. mCherry was used as loading control and detected with anti-RFP antibody. GAPDH was used as a loading control. **(D)** Flow cytometry plots of the COX4I1- and COX4I2 reporter HEK293T cells as in (C). **(E)** MARCHF6 co-precipitates specifically with COX4I1 precursor but not the mature proteins. Extracts of the indicated cell lines were subjected to immunoprecipitation with anti-HA beads and eluted proteins were analysed by SDS-PAGE and immunoblotting. COX4I1 was detected with anti-FLAG antibodies. MARCHF6 variants were detected with anti-HA antibodies. The abundant ER membrane protein Climp-63 was used as specificity control and detected with an anti-CLIMP63 antibody. **(F)** Flow cytometry analysis of HEK293T cells expressing the indicated COX4 WT and chimeras as part of the split-moxVenus reporter system. The various cell lines were subject to MARCHF6 sgRNAs as indicated.

Despite their distinct fates when mislocalized to the ER, COX4I1 and COX4I2 share a similar structural organization (Figure S2B-D). To gain insight into the features of mislocalized COX4I1 recognized by MARCHF6, we generated various COX4I1- COX4I2 chimeric proteins fused to moxV(C). The fate of these chimeras was assessed in cells with constitutive expression of ER-moxV(N)-K (Figure 4F). Swapping the MTS or the C-terminal regions (CTR) did not appreciably alter the behaviour of the chimeras. In contrast, inserting the N-terminal domain (NTD) of COX4I1 was sufficient to destabilise an otherwise stable COX4I2 backbone, and this destabilisation required MARCHF6 (Figure 4F). Thus, the NTD of COX4I1 encodes a transferable, MARCHF6-dependent degron.

### Additional ERAD factors contribute for the clearance ER mistargeted OXPHOS proteins

The turnover of mislocalized COX4I1 and UQCR10 was strongly diminished but not completely blocked in *MARCHF6* KO cells (Figure 3C and 4C). This observation suggested that additional pathways contribute to the clearance of mistargeted mitochondrial proteins from the ER.

To identify factors working in parallel with MARCHF6 in the clearance of mistargeted mitochondrial proteins, we performed a genome-wide screen analogous to that described above in *MARCHF6* KO cells. Among the top hits were components of other ERAD branches, including the ubiquitin ligase SYVN1/HRD1, its binding partners SEL1L and DERL2 as well as the ubiquitin ligase RNF145 (Figure 5A and S3A). Independent sgRNAs targeting a selected subset of hits were used to validate the screen and their effect were analyzed in the presence and absence of *MARCHF6* (Figure 5B). In cells that retained MARCHF6, loss of HRD1 or other ERAD factors had no detectable effect on the levels of COX4I1. Instead, the contribution of these factors became apparent in MARCHF6 KO cells (Figure 5B).

**Figure 5.**
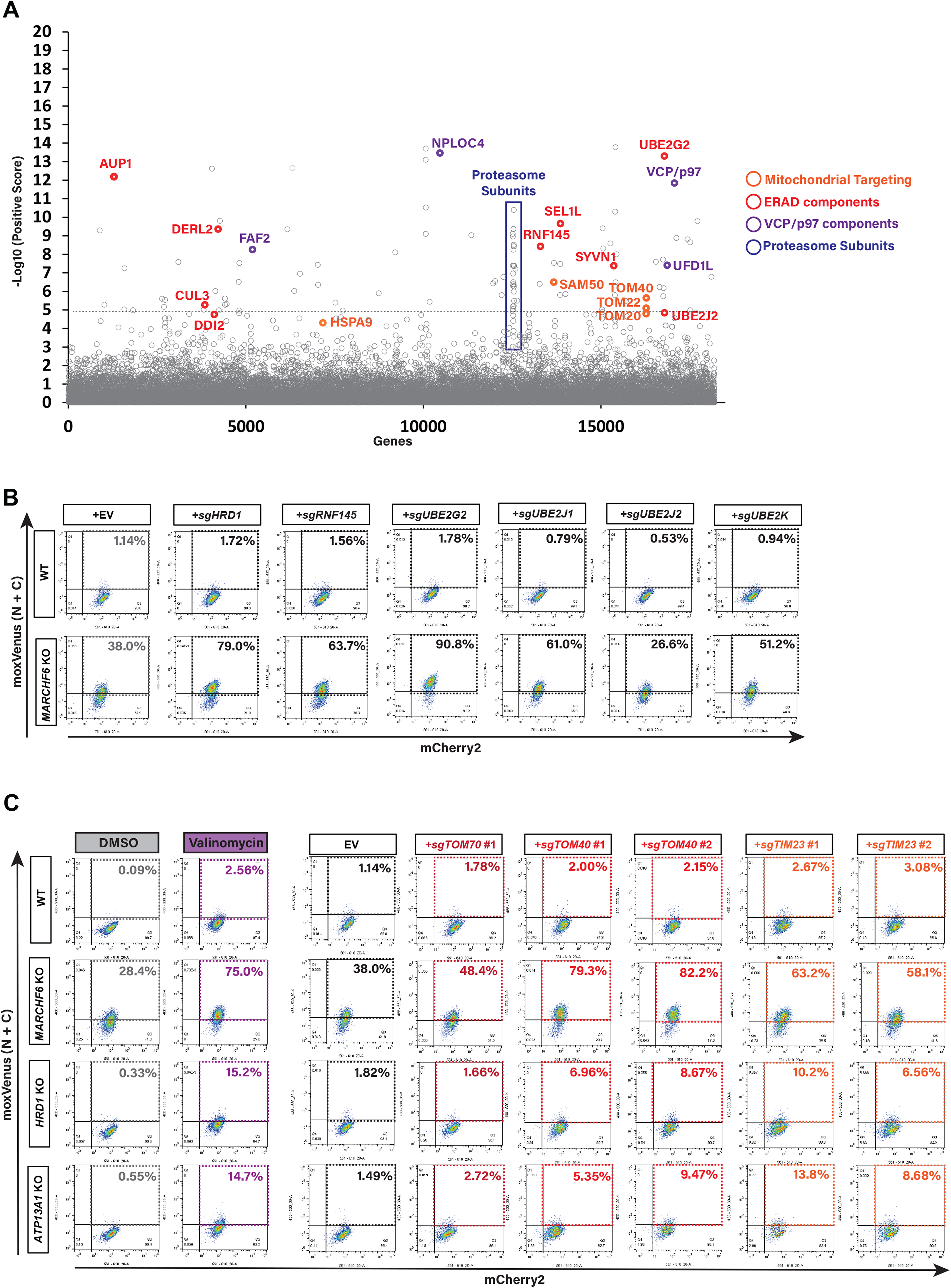
Degradation of ER mistargeted COX4I1 involves partly redundant ERAD factors. **(A)** Significance score of the genes analyzed in the screen calculated by the MAGeCK algorithm. The x axis represents the genes in alphabetical order. The y axis shows the −log(αRRA) significance value. The −log(αRRA) cutoff was arbitrarily set at 9 (dashed line). Significantly enriched genes are annotated. MARCHF6 KO cells expressing the COX4I1 reporter were transduced with the TKOv3 sgRNA library and the population with high levels of moxVenus fluorescence was isolated by flow cytometry 8 days post transduction. **(B)** Parental or MARCHF6 KO cells expressing the COX4I1 split-moxVenus reporter were depleted of the indicated ERAD factors and analysed by flow cytometry. The percentage of moxVenus-positive cells is indicated in each case. **(C)** Parental cells or the indicated KO cells expressing the COX4I1 split-moxVenus reporter were analysed by flow cytometry following the indicated treatment or sgRNA transfection. The percentage of moxVenus-positive cells is indicated in each case.

We next triggered import stress by depleting several components of the mitochondrial import machinery. In addition to the ERAD factors identified by our genetic screens, we also examined the contribution of ATP13A1, a conserved P5 ATPase recently shown to promote the extraction of mistargeted and misinserted tail-anchored (TA) membrane proteins from the ER (Dederer et al., 2019; McKenna et al., 2020, 2022). This analysis confirmed that the clearance of mislocalized COX4I1 depends primarily on MARCHF6, both in the absence and presence of diverse levels of mitochondrial import stress (Figure 5C). By comparison, the contribution of HRD1 or ATP13A1 was more modest and only measurable under conditions of severe mitochondrial stress, possibly because the MARCHF6-dependent pathway becomes saturated.

## Discussion

In this study, we establish an integrated approach to systematically investigate the fate of mitochondrial proteins that become mistargeted to the endoplasmic reticulum (ER). By combining ER-targeted proximity labelling, a split-fluorescence reporter and genome-wide CRISPR screening, we define both the scope of mitochondrial protein mistargeting and the mechanisms that govern their clearance.

Our data show that mitochondrial import stress leads to a broad but selective rerouting of mitochondrial proteins to the ER, with a strong enrichment for oxidative phosphorylation (OXPHOS) components (Figure 1C-E). These proteins are highly hydrophobic and frequently contain transmembrane domains, features that likely increase their propensity to engage ER targeting pathways when mitochondrial import is compromised. This observation is consistent with emerging evidence that protein targeting is highly dynamic and influenced by nascent chain properties (Luo et al., 2025). For example, ribosome-associated factors such as NAC antagonize signal recognition particle (SRP)-mediated ER targeting, thereby preventing mistargeting of non-secretory proteins cotranslationally (Jomaa et al., 2022; Gamerdinger et al., 2026). Under conditions of mitochondrial import stress, this balance may be perturbed, allowing mitochondrial precursors—especially hydrophobic ones—to be aberrantly captured by ER-targeting pathways.

Although the precise route by which mitochondrial proteins access the ER remains to be defined, our data are most consistent with a model in which hydrophobic precursor proteins are aberrantly engaged by ER-targeting pathways under conditions of import stress. Increased SRP engagement or reduced NAC-mediated shielding are plausible mechanisms, given the known roles of these factors in maintaining targeting fidelity. Contributions from alternative pathways, such as the GET system (Vitali et al., 2018), ER-SURF (Hansen et al., 2018) or passive overflow, cannot be excluded and should be tested in future studies.

An interesting observation is that mistargeted mitochondrial proteins may not be handled uniformly but instead segregate into distinct behaviors. While some proteins (e.g., COX4I2, COX16 and COA3) appear to remain stable in the ER environment, others (e.g., COX4I1 and UQCR10) are efficiently degraded by ERAD. A notable example concerns COX4I1 and COX4I2 which share similar amino acid sequences and structural features but their stability within the ER differs strikingly (Figure 4). Mechanistically, our data indicates that in this case substrate recognition is not dictated by mitochondrial targeting sequences per se but instead by determinants within COX4I1 N-terminal domain (NTD), which are sufficient to impose MARCHF6-dependent degradation on the otherwise stable COX4I2 backbone. The precise feature recognized within this region remains to be defined. These findings are consistent with the idea that defects in topology and membrane insertion may act as key signals for ERAD engagement, consistent with broader models in which ER quality control senses aberrant membrane protein features (McKenna and Shao, 2023; Sergejevs and Carvalho, 2025).

Our genetic screens identify the ER-resident ubiquitin ligase MARCHF6 as a central component of the quality control machinery responsible for clearing mistargeted mitochondrial proteins. MARCHF6 loss leads to accumulation of precursor forms such as COX4I1 and UQCR10, demonstrating its primary role in ERAD of these substrates. Additional ERAD branches, including HRD1 and RNF145, contribute in parallel, particularly under conditions of elevated stress. This suggests that for the clearance of OXPHOS proteins, multiple ERAD pathways operate in parallel, with MARCHF6 acting as dominant ligase and other ERAD branches providing backup capacity when substrate load increases.

Together, these findings support a model in which mitochondrial import stress generates a pool of precursor proteins some of which are redirected to the ER, where they are triaged (Koch et al., 2024; Hansen et al., 2018; Kroczek et al., 2026; Xiao et al., 2021). Proteins compatible with the ER environment can persist even if nonfunctional, whereas aberrant substrates are recognized and eliminated by ERAD pathways (Gamerdinger et al., 2015; Laborenz et al., 2021; Knöringer et al., 2023).

While our study establishes a framework for understanding ER-mediated handling of mistargeted mitochondrial proteins, several questions remain. In particular, the molecular routes that mediate ER targeting and the determinants of substrate classification require further investigation. In addition, although our systems allow controlled analysis of mistargeting, future work will be needed to assess how these processes operate under physiological and pathological conditions. Overall, our findings reveal that the ER plays an active and selective role in maintaining proteome integrity during mitochondrial stress, acting as a critical interface between protein targeting fidelity and cellular quality control.

## Materials and methods

### Growth conditions

Flp-In^TM^, TREx^®^ HEK cells were obtained from Invitrogen (#R75007, Thermo Fischer Scientific). Flp-In^TM^, TREx^®^ HEK clones were established using manufacturer’s guidelines. 293T Lenti-X virus packaging cells (#Z2180N/#632180) were obtained from Takara Clontech. All cells were grown at 37°C, 5% CO_2_ in DMEM medium (Sigma-Aldrich) supplemented with L-glutamine (2 mM; Gibco), penicillin–streptomycin (10 units/ml; Gibco), and 10% Fetal Calf Serum (FCS) (Gibco). Cells were routinely checked for mycoplasma contamination using in-house MycoAlert® Mycoplasma Detection Kit (LT07-318; Lonza) and confirmed to be negative.

### Lentivirus production

For individual gene infections using lentivirus, virus was produced in 6-well plates using TransIT LT1 (#MIR2304, TaKaRa) and second-generation packaging vectors according to standard lentiviral production protocols.

### Gene deletion by transfection of sgRNA-Cas9 plasmid, RP418

Flp-In^TM^, TREx^®^ HEK cells were seeded at 2.5*10^5 cells/well in 24 well plates and cultured in CO_2_ incubator for 12-16 hrs. 500 ng of RP418 plasmids were transfected by TransIT^®^-LT1 transfection reagent (#MIR2304, TaKaRa). Culture media were exchanged to DMEM complete medium containing 5 μg/mL puromycin and cells were cultured for 2 days. Cells were recovered by exchanging culture medium to DMEM complete medium without puromycin for 2-3 days and analyzed by immunoblotting, immunofluorescence and FACS analysis. With the exception of TKOv3 library, all other sgRNAs used in this study have been fully validated in this or prior studies (van de Weijer et al., 2020).

### ER-APEX2-K biotinylation pulldown

1.0*10^7 cells of Flp-In T-REx 293 stably expressing BiPss-APEX2-KDEL cells were seeded onto collagen typeI-C coated 10-cm dishes, and cultures for 24 hours. BiPss-APEX2-KDEL HEK293T cells were treated with mistargeting cocktail, 1 μM Valinomycin, 1 μM Bortezomib and 250 nM ISRIB, for 18 hrs. BiPss-APEX2-KDEL HEK293T cells were incubated with 500 nM Biotin-phenol for 30 min and then treated with 10 mM H_2_O_2_ for 30 sec. Cells were washed with ice-cold 1xPBS(-) containing 10 mM sodium ascorbate, 10 mM sodium azide, 5 mM Trolox three times and collected to 50-mL Falcon tube and spined down at 500 r.c.f. for 5 min. Cells were lysed by adding RIPA buffer containing 10 mM sodium ascorbate, 10 mM sodium azide, 5 mM Trolox on ice for 5 min. Biotin modified proteome were pulled down following the recent APEX2 methods (Hung et al., 2016).

### Cycloheximide chase

One day prior, 1.2x10^5 cells were seeded in a 24-well plate, pre-coated with Cellmatrix® Type I -C (#Collagen Type I-C, Nitta Gelatin), and protein expression was induced with 100 ng/mL doxycycline for 16 hours. The following day, cells were incubated with cycloheximide (50 μg/mL) for the time indicated, after which the media was aspirated and the cells were directly lysed in 1x Laemmli sample buffer containing Benzonase (E1014; Merck), cOmplete™ protease inhibitor cocktail and 100 mM DTT. Lysates were incubated on a ThermoMixer for 20 minutes at 37 oC, after which denatured samples were either frozen or used directly for immunoblotting.

### Genome-wide CRISPR-Cas9 screen

The TKOv3 CRISPR-Cas9 library was a gift from Jason Moffat (Addgene #90294). The sgRNA library and 2nd generation lentiviral packaging plasmids (psPAX2 and pMD2.G) were co-transfected into 293T cells. After 48 and 72 hours, lentivirus was harvested, concentrated by ultra-centrifugation, and infected into 125*10^6 Flp-In T-REx 293 expressing COX4I1-3Flag-moxVenus(C)-KDEL/BiPss-mCherry2-moxVenus(N)-KDEL cells at an MOI of 0.3 to achieve a 250-fold coverage of the library after selection. Cells were grown for 48 h and then selected with 2 μg/mL puromycin (#A1113803, GIBCO) for 72 hours. Cells were then split into two technical replicates. 24 hours prior to sorting, 100 ng/mL doxycycline was added to the cell medium to induce the expression of the COX4i1 reporter. 2% of the brightest Venus (Venus^high^) cells from 25*10^6 cells were collected using a BD FACSAria3 and a Beckman Coulter MoFlo Astrios. Genomic DNA was extracted from each cell population using a QIAGEN BloodMaxi kit (for reference samples) or BloodMini kit (for sorted Venus^high^ samples) according to the manufacturer’s protocol. SgRNAs were PCR amplified from the entire isolated genomic DNA using NEBNext® Ultra II Q5® Master Mix (#M0544X, NEB) and the primers v2.1-F1 and v2.1-R1, according to the TKOv3 protocol. PCR reactions were pooled again, after which a second PCR was performed to attach indices and sequencing adapters using the primers i5 and i7. The PCR reaction was loaded onto a 2% agarose gel, the 200bp band excised and purified using a GeneJet PCR Purification kit (#K0701, Thermo Fischer Scientific). Libraries were analyzed by deep sequencing on an Illumina HiSeq4000. Gene rankings were generated using the MAGeCK algorithm

### Co-Immunoprecipitation

3*10^6 HEK293TF cells were seeded on 10-cm dishes, pre-coated with poly-L-lysine, in DMEM. 24 hours after seeding, DMEM was changed to media containing doxycycline for construct expression and, where stated, mistargeting cocktail (1 μM Valinomycin, 400 nM ISRIB, 1 μM Bortezomib) for 16 hours. Cells were treated with 2.5 μM CB-5083 4h prior to harvesting. Cells were washed with TBS once (50 mM Tris-HCl pH 7.5, 150 mM NaCl) and harvested in 1% Decyl Maltose Neopentyl Glycol (DMNG) (#DG322, Anatrace) lysis buffer (50 mM Tris-HCl pH 7.5, 150 mM NaCl) containing cOmplete protease inhibitor cocktail (Roche) by scraping from the dishes on ice. Cell suspension was solubilized with head-over-head rotation for 2h at 4℃, followed by centrifugation (20,000 x g) for 20 minutes at 4℃. Recovered post-nuclear supernatant was incubated for 2 hours with 30 μL pre-equilibrated anti-HA magnetic beads (#SAE0197, Sigma-Aldrich) with head-over-head rotation at 4°C. After three 10 minutes washes in 0.1% DMNG washing buffer (50 mM Tris-HCl pH 7.5, 150 mM NaCl), immunoprecipitated proteins were eluted from magnetic beads with 1x Laemmli sample buffer for 10 minutes at 65℃ and transferred to a fresh tube using magnetic rack and supplemented with 100 mM DTT. Denatured material was either stored in -20℃ or used directly for immunoblotting as described.

### Immunofluorescence

HEK293TF cells (5.0*10^4) were seeded onto round coverslips in a 12-well tissue culture-treated plate pre-coated with Poly-L-Lysine. 24 hours later, media was replaced with complete DMEM supplemented with either Doxycycline alone or Doxycycline with 2.5 μM Bortezomib and the cells were incubated for further 14 hours. Cell culture media was aspirated, and the cells were washed with PBS once. Following this, cells were fixed by adding 1mL of 4% methanol-free Paraformaldehyde (PFA) in PBS for 15 minutes at 37℃. PFA was removed and fixed cells were washed three times with PBS and incubated in 1 mL blocking buffer (0.1% Saponin, 1% BSA in PBS) for 15 minutes at room temperature. Following two washes with PBS, permeabilized cells were incubated with primary antibody in blocking buffer for 1 hour in the dark, by placing the coverslips onto pre-spotted antibody aliquots on parafilm underneath custom aluminium foil-covered moisturising chamber. Cover slips were then transferred to fresh 12-well plate and washed with blocking buffer three times. Fluorophore-conjugated secondary antibody incubation was done in a similar way to primary antibody incubation. Cover slips were washed with PBS and incubated in 1 mL of DAPI-containing PBS for 5 minutes at room temperature, followed by three 5-minute washes with PBS. Coverslips were then mounted on microscopy glass slides in a non-hardening mounting media (H-1900; Vector Laboratories) and sealed with a nail polish. Samples were imaged immediately after full solidification of mounting media or kept at 4℃ until further processing.

### Fluorescence Microscopy

Fixed cells on slides were imaged at room temperature using Zeiss Axio Observer Z1 equipped with a complementary metal-oxide semiconductor (CMOS) camera (Hamamatsu ORCA-Flash4.0), controlled by 3i Slidebook 6.0 software. The system was equipped with Plan-Apochromat ×63/1.4-NA oil lens, with an immersion oil (Immersol W 2010, Carl Zeiss; refractive index of 1.518). The resulting images were exported as .TIFF files.

### Mass Spectrometry

Immunoprecipitated samples on beads were re-suspended in 25 μL 1x SDS sample buffer (5% SDS, 50mM TEAB, pH 7.55). Disulfide bonds were reduced using 20 mM TCEP for 15 mins at 47℃. Following this, cooled samples were alkylated using 20mM CAA in the dark for 15 minutes and 10% volume of 12% phosphoric acid was added to acidify the samples. S-trap binding buffer (90% methanol in 100 mM TEAB, pH 7.5) was added to acidified, denatured samples to a final volume of 190uL and the resulting solution was loaded onto S-Trap micro spin columns, with a maximum of 150 μL of sample per load. Loaded spin columns were centrifuged at 4000g for 1 minute and this step was repeated until entire sample is loaded onto a spin column. S-Trap columns were washed 5x with S-trap binding buffer (90% methanol in 100 mM TEAB, pH 7.5) and the columns were moved onto 2 mL low protein binding Eppendorf tubes. To each S-trap column, 25 μL of digestion solution (50 mM TEAB, pH 8.0), containing 2 μg of Trypsin/LysC mix (Promega) was added and loosely capped columns were incubated for 3 hours at 47℃ on a ThermoMixer. Peptides were eluted with 30 μL of 50 mM TEAB, followed by 30 μL of 0.2% formic acid and 40 μL of 50% acetonitrile in 0.2% formic acid. Peptides were dried for 4 hours at 37℃ in a vacuum centrifuge and samples were stored at -80℃ until further analysis.

Mass spectrometric identification and quantification were performed on an Orbitrap Fusion Tribid mass spectrometer. Protein identification and quantification were performed using MaxQuant (version 1.6.10.43). Search was conducted using the Uniprot-SwissProt Homo sapiens database (containing 42,371 database entries with isoforms, retrieved on 2021/02/24). Identification were filtered at a 1% false-discovery rate (FDR) at the protein level, accepting a minimum peptide length of 7 amino acids. Quantification of identified proteins referred to razor and unique peptides required a minimum ratio count of 2.

### Flow Cytometry

Cells were trypsinised, washed once with ice-cold PBS and re-suspended in FACS buffer (1 mM EDTA, 2 % FBS in PBS). Cells were then assessed for expression of target constructs by fluorescence using a BD LSRFortessa X-20 and data was processed using FlowJo. At least 10,000 cells per sample were analysed, gating on the main population in the forward scatter/side scatter (FSC/SSC) plot.

### Quantification and Statistical analysis

Western blot data were acquired and analysed using Image lab software (Bio-Rad) and representative images of at least three independent experiments are shown.

## Data availability

The raw reads dataset generated during this study is available at NCBI SRA database with the reference PRJNA1507838.

The mass spectrometry proteomics data have been deposited to the ProteomeXchange Consortium via the PRIDE partner repository with the dataset identifier PXD081401.

## Acknowledgments

We thank Michael van de Weijer and Logesvaran Krshnan for help with CRISPR screen analysis, L. Witty for help with high-throughput sequencing, M. Maj and L. Eriksen for help with flow cytometry. P.C. was supported by an investigator award from The Wellcome Trust (202642/Z/16/Z) and an ERC consolidator grant (GA 817708).

## Author contributions

Y.T. and P.C. designed the study. Y.T. performed most of the experiments with the help of N.S., and M.R.; N.S. performed the HA-pulldown assay; M.R. helped with the establishing split moxVenus assay. M.E.D. and M.T. performed the mass spectrometry analysis. Y.T. and P.C. analyzed the data with the help of all the authors. P.C. and Y.T. wrote the manuscript with input from all the authors.

## Legends to the Supplemental Figures

**Supplemental Figure 1. Characterization of factors involved in quality of mistargeted COX4I1.**

**(A)** Flow cytometry plots of HEK293T cells expressing indicated reporter with moxVenus(C) and ER-luminal facing [BiPss-mCherry2-moxVenus(N)-KDEL (ER-moxV(N)-KDEL)] or ER-cytosolic facing acceptors [mCherry2-moxVenus(N)-CytB5 (moxV(N)-CytB5)].

**(B)** Significance score of the genes analyzed in the screen calculated by the MAGeCK algorithm. The x axis represents the genes in alphabetical order. The y axis shows the −log(αRRA) significance value. The −log(αRRA) cutoff was arbitrarily set at 9 (dashed line). Significantly enriched genes are annotated. COX4I1 reporter expressing cells were transduced with the TKOv3 sgRNA library and the cell population with high moxVenus fluorescence levels was isolated by flow cytometry 15 days post transduction.

**Supplemental Figure 2. Comparison of COX4I1 and COX4I2 proteins.**

**(A)** Extracts of Parental and indicated KO cells expressing the COX4I1 reporter were analysed by SDS-PAGE and immunoblotting.

**(B)** Schematic representation of *Homo Sapiens* COX4I1 and COX4I2 domains (MTS, N-terminus disordered region (NTD), transmembrane domain (TMD) and C-terminus disordered region (CTR)).

**(C)** Alignment of amino acid sequence between *Homo Sapiens* COX4I1 and COX4I2 protein.

**(D)** Comparison of AlphaFold 3 models of *Homo Sapiens* COX4I1 and COX4I2 proteins.

**Supplemental Figure 3. Identification of additional factors involved in the degradation of mistargeted COX4I1.**

**(A)** Significance score of the genes analyzed in the screen calculated by the MAGeCK algorithm. The x axis represents the genes in alphabetical order. The y axis shows the −log(αRRA) significance value. The −log(αRRA) cutoff was arbitrarily set at 9 (dashed line). Significantly enriched genes are annotated. MARCHF6 KO cells expressing the COX4I1 reporter were transduced with the TKOv3 sgRNA library and the population with high levels of moxVenus fluorescence was isolated by flow cytometry 15 days post transduction.

## References

Boos, F., L. Krämer, C. Groh, F. Jung, P. Haberkant, F. Stein, F. Wollweber, A. Gackstatter, E. Zöller, M. van der Laan, M.M. Savitski, V. Benes, and J.M. Herrmann. 2019. Mitochondrial protein-induced stress triggers a global adaptive transcriptional programme. Nat. Cell Biol. 21:442–451. doi:10.1038/s41556-019-0294-5.

den Brave, F., U. Schulte, B. Fakler, N. Pfanner, and T. Becker. 2024. Mitochondrial complexome and import network. Trends Cell Biol. 34:578–594. doi:10.1016/j.tcb.2023.10.004.

Bykov, Y.S., T. Flohr, F. Boos, N. Zung, J.M. Herrmann, and M. Schuldiner. 2022. Widespread use of unconventional targeting signals in mitochondrial ribosome proteins. EMBO J. 41:e109519. doi:10.15252/embj.2021109519.

Bykov, Y.S., D. Rapaport, J.M. Herrmann, and M. Schuldiner. 2020. Cytosolic events in the biogenesis of mitochondrial proteins. Trends Biochem. Sci. 45:650–667. doi:10.1016/j.tibs.2020.04.001.

Christianson, J.C., and P. Carvalho. 2022. Order through destruction: how ER-associated protein degradation contributes to organelle homeostasis. EMBO J. 41:e109845. doi:10.15252/embj.2021109845.

Christianson, J.C., E. Jarosch, and T. Sommer. 2023. Mechanisms of substrate processing during ER-associated protein degradation. Nat. Rev. Mol. Cell Biol. 24:777–796. doi:10.1038/s41580-023-00633-8.

Costantini, L.M., M. Baloban, M.L. Markwardt, M.A. Rizzo, F. Guo, V.V. Verkhusha, and E.L. Snapp. 2015. A palette of fluorescent proteins optimized for diverse cellular environments. Nat. Commun. 6:7670. doi:10.1038/ncomms8670.

Dederer, V., A. Khmelinskii, A.G. Huhn, V. Okreglak, M. Knop, and M.K. Lemberg. 2019. Cooperation of mitochondrial and ER factors in quality control of tail-anchored proteins. eLife. 8. doi:10.7554/eLife.45506.

Endo, T., and N. Wiedemann. 2025. Molecular machineries and pathways of mitochondrial protein transport. Nat. Rev. Mol. Cell Biol. 26:848–867. doi:10.1038/s41580-025-00865-w.

Friedman, J.R., M. Kannan, A. Toulmay, C.H. Jan, J.S. Weissman, W.A. Prinz, and J. Nunnari. 2018. Lipid homeostasis is maintained by dual targeting of the mitochondrial PE biosynthesis enzyme to the ER. Dev. Cell. 44:261–270.e6. doi:10.1016/j.devcel.2017.11.023.

Gamerdinger, M., N. Burg, and E. Deuerling. 2026. Ribosome-NAC collaboration: A regulatory platform for cotranslational chaperones, enzymes, and targeting factors. Mol. Cell. 86:491–502. doi:10.1016/j.molcel.2025.12.031.

Gamerdinger, M., M.A. Hanebuth, T. Frickey, and E. Deuerling. 2015. The principle of antagonism ensures protein targeting specificity at the endoplasmic reticulum. Science. 348:201–207. doi:10.1126/science.aaa5335.

Guna, A., and R.S. Hegde. 2018. Transmembrane Domain Recognition during Membrane Protein Biogenesis and Quality Control. Curr. Biol. 28:R498–R511. doi:10.1016/j.cub.2018.02.004.

Hansen, K.G., N. Aviram, J. Laborenz, C. Bibi, M. Meyer, A. Spang, M. Schuldiner, and J.M. Herrmann. 2018. An ER surface retrieval pathway safeguards the import of mitochondrial membrane proteins in yeast. Science. 361:1118–1122. doi:10.1126/science.aar8174.

Hart, T., A.H.Y. Tong, K. Chan, J. Van Leeuwen, A. Seetharaman, M. Aregger, M. Chandrashekhar, N. Hustedt, S. Seth, A. Noonan, A. Habsid, O. Sizova, L. Nedyalkova, R. Climie, L. Tworzyanski, K. Lawson, M.A. Sartori, S. Alibeh, D. Tieu, S. Masud, P. Mero, A. Weiss, K.R. Brown, M. Usaj, M. Billmann, M. Rahman, M. Constanzo, C.L. Myers, B.J. Andrews, C. Boone, D. Durocher, and J. Moffat. 2017. Evaluation and Design of Genome-Wide CRISPR/SpCas9 Knockout Screens. G3 (Bethesda). 7:2719–2727. doi:10.1534/g3.117.041277.

Hegde, R.S., and R.J. Keenan. 2022. The mechanisms of integral membrane protein biogenesis. Nat. Rev. Mol. Cell Biol. 23:107–124. doi:10.1038/s41580-021-00413-2.

Hung, V., N.D. Udeshi, S.S. Lam, K.H. Loh, K.J. Cox, K. Pedram, S.A. Carr, and A.Y. Ting. 2016. Spatially resolved proteomic mapping in living cells with the engineered peroxidase APEX2. Nat. Protoc. 11:456–475. doi:10.1038/nprot.2016.018.

Jomaa, A., M. Gamerdinger, H.-H. Hsieh, A. Wallisch, V. Chandrasekaran, Z. Ulusoy, A. Scaiola, R.S. Hegde, S.-O. Shan, N. Ban, and E. Deuerling. 2022. Mechanism of signal sequence handover from NAC to SRP on ribosomes during ER-protein targeting. Science. 375:839–844. doi:10.1126/science.abl6459.

Knöringer, K., C. Groh, L. Krämer, K.C. Stein, K.G. Hansen, J. Zimmermann, B. Morgan, J.M. Herrmann, J. Frydman, and F. Boos. 2023. The unfolded protein response of the endoplasmic reticulum supports mitochondrial biogenesis by buffering nonimported proteins. Mol. Biol. Cell. 34:ar95. doi:10.1091/mbc.E23-05-0205.

Koch, C., S. Lenhard, M. Räschle, C. Prescianotto-Baschong, A. Spang, and J.M. Herrmann. 2024. The ER-SURF pathway uses ER-mitochondria contact sites for protein targeting to mitochondria. EMBO Rep. 25:2071–2096. doi:10.1038/s44319-024-00113-w.

Krämer, L., N. Dalheimer, M. Räschle, Z. Storchová, J. Pielage, F. Boos, and J.M. Herrmann. 2023. MitoStores: chaperone-controlled protein granules store mitochondrial precursors in the cytosol. EMBO J. 42:e112309. doi:10.15252/embj.2022112309.

Kroczek, L., H. Nolte, Y. Lasarzewski, I. Agrawal, T. Molinié, D. Curbelo Piñero, K. Lemke, E. Rugarli, and T. Langer. 2026. Stress adaptation of mitochondrial protein import by OMA1-mediated degradation of DNAJC15. Nat. Struct. Mol. Biol. 33:499–511. doi:10.1038/s41594-026-01756-0.

Krshnan, L., M.L. van de Weijer, and P. Carvalho. 2022. Endoplasmic Reticulum-Associated Protein Degradation. Cold Spring Harb. Perspect. Biol. 14. doi:10.1101/cshperspect.a041247.

Laborenz, J., Y.S. Bykov, K. Knöringer, M. Räschle, S. Filker, C. Prescianotto-Baschong, A. Spang, T. Tatsuta, T. Langer, Z. Storchová, M. Schuldiner, and J.M. Herrmann. 2021. The ER protein Ema19 facilitates the degradation of nonimported mitochondrial precursor proteins. Mol. Biol. Cell. 32:664–674. doi:10.1091/mbc.E20-11-0748.

Lee, J.H., L. Rabl, M. Gamerdinger, V. Goyal, K.M. Khakzar, N.M. Barbosa, J. Abramovich, F. Morales-Polanco, A.-K. Köhler, E. Samatova, M.V. Rodnina, E. Deuerling, and J. Frydman. 2026. NAC controls nascent chain fate through tunnel sensing and chaperone action. Nature. 652:230–239. doi:10.1038/s41586-025-10058-2.

Liu, Y.J., J. Sulc, and J. Auwerx. 2025. Mitochondrial genetics, signalling and stress responses. Nat. Cell Biol. 27:393–407. doi:10.1038/s41556-025-01625-w.

Li, W., H. Xu, T. Xiao, L. Cong, M.I. Love, F. Zhang, R.A. Irizarry, J.S. Liu, M. Brown, and X.S. Liu. 2014. MAGeCK enables robust identification of essential genes from genome-scale CRISPR/Cas9 knockout screens. Genome Biol. 15:554. doi:10.1186/s13059-014-0554-4.

Luo, J., S. Khandwala, J. Hu, S.-Y. Lee, K.L. Hickey, Z.G. Levine, J.W. Harper, A.Y. Ting, and J.S. Weissman. 2025. Proximity-specific ribosome profiling reveals the logic of localized mitochondrial translation. Cell. 188:5589–5604.e17. doi:10.1016/j.cell.2025.08.002.

Matsumoto, S., K. Nakatsukasa, C. Kakuta, Y. Tamura, M. Esaki, and T. Endo. 2019. Msp1 Clears Mistargeted Proteins by Facilitating Their Transfer from Mitochondria to the ER. Mol. Cell. 76:191–205.e10. doi:10.1016/j.molcel.2019.07.006.

McKenna, M.J., B.M. Adams, V. Chu, J.A. Paulo, and S. Shao. 2022. ATP13A1 prevents ERAD of folding-competent mislocalized and misoriented proteins. Mol. Cell. 82:4277–4289.e10. doi:10.1016/j.molcel.2022.09.035.

McKenna, M.J., and S. Shao. 2023. The endoplasmic reticulum and the fidelity of nascent protein localization. Cold Spring Harb. Perspect. Biol. 15. doi:10.1101/cshperspect.a041249.

McKenna, M.J., S.I. Sim, A. Ordureau, L. Wei, J.W. Harper, S. Shao, and E. Park. 2020. The endoplasmic reticulum P5A-ATPase is a transmembrane helix dislocase. Science. 369. doi:10.1126/science.abc5809.

Michaelis, J.B., M.E. Brunstein, S. Bozkurt, L. Alves, M. Wegner, M. Kaulich, C. Pohl, and C. Münch. 2022. Protein import motor complex reacts to mitochondrial misfolding by reducing protein import and activating mitophagy. Nat. Commun. 13:5164. doi:10.1038/s41467-022-32564-x.

Morizono, M.A., K.L. McGuire, N.I. Birouty, and M.A. Herzik. 2024. Structural insights into GrpEL1-mediated nucleotide and substrate release of human mitochondrial Hsp70. BioRxiv. doi:10.1101/2024.05.10.593630.

Muthukumar, G., T.A. Stevens, A.J. Inglis, T.K. Esantsi, R.A. Saunders, F. Schulte, R.M. Voorhees, A. Guna, and J.S. Weissman. 2024. Triaging of α-helical proteins to the mitochondrial outer membrane by distinct chaperone machinery based on substrate topology. Mol. Cell. 84:1101–1119.e9. doi:10.1016/j.molcel.2024.01.028.

Nowicka, U., P. Chroscicki, K. Stroobants, M. Sladowska, M. Turek, B. Uszczynska-Ratajczak, R. Kundra, T. Goral, M. Perni, C.M. Dobson, M. Vendruscolo, and A. Chacinska. 2021. Cytosolic aggregation of mitochondrial proteins disrupts cellular homeostasis by stimulating the aggregation of other proteins. eLife. 10. doi:10.7554/eLife.65484.

Pfanner, N., F. den Brave, and T. Becker. 2025. Mitochondrial protein import stress. Nat. Cell Biol. 27:188–201. doi:10.1038/s41556-024-01590-w.

Rackham, O., and A. Filipovska. 2022. Organization and expression of the mammalian mitochondrial genome. Nat. Rev. Genet. 23:606–623. doi:10.1038/s41576-022-00480-x.

Sergejevs, N., and P. Carvalho. 2025. Mechanisms of transmembrane domain recognition during endoplasmic reticulum quality control. Curr. Opin. Cell Biol. 96:102580. doi:10.1016/j.ceb.2025.102580.

Shakya, V.P., W.A. Barbeau, T. Xiao, C.S. Knutson, M.H. Schuler, and A.L. Hughes. 2021. A nuclear-based quality control pathway for non-imported mitochondrial proteins. eLife. 10. doi:10.7554/eLife.61230.

Song, J., J.M. Herrmann, and T. Becker. 2021. Quality control of the mitochondrial proteome. Nat. Rev. Mol. Cell Biol. 22:54–70. doi:10.1038/s41580-020-00300-2.

Stefanovic-Barrett, S., A.S. Dickson, S.P. Burr, J.C. Williamson, I.T. Lobb, D.J. van den Boomen, P.J. Lehner, and J.A. Nathan. 2018. MARCH6 and TRC8 facilitate the quality control of cytosolic and tail-anchored proteins. EMBO Rep. 19. doi:10.15252/embr.201745603.

Vercellino, I., and L.A. Sazanov. 2022. The assembly, regulation and function of the mitochondrial respiratory chain. Nat. Rev. Mol. Cell Biol. 23:141–161. doi:10.1038/s41580-021-00415-0.

Vitali, D.G., M. Sinzel, E.P. Bulthuis, A. Kolb, S. Zabel, D.G. Mehlhorn, B. Figueiredo Costa, Á. Farkas, A. Clancy, M. Schuldiner, C. Grefen, B. Schwappach, N. Borgese, and D. Rapaport. 2018. The GET pathway can increase the risk of mitochondrial outer membrane proteins to be mistargeted to the ER. J. Cell Sci. 131. doi:10.1242/jcs.211110.

van de Weijer, M.L., L. Krshnan, S. Liberatori, E.N. Guerrero, J. Robson-Tull, L. Hahn, R.J. Lebbink, E.J.H.J. Wiertz, R. Fischer, D. Ebner, and P. Carvalho. 2020. Quality control of ER membrane proteins by the rnf185/membralin ubiquitin ligase complex. Mol. Cell. 79:768–781.e7. doi:10.1016/j.molcel.2020.07.009.

Wrobel, L., U. Topf, P. Bragoszewski, S. Wiese, M.E. Sztolsztener, S. Oeljeklaus, A. Varabyova, M. Lirski, P. Chroscicki, S. Mroczek, E. Januszewicz, A. Dziembowski, M. Koblowska, B. Warscheid, and A. Chacinska. 2015. Mistargeted mitochondrial proteins activate a proteostatic response in the cytosol. Nature. 524:485–488. doi:10.1038/nature14951.

Xiao, T., V.P. Shakya, and A.L. Hughes. 2021. ER targeting of non-imported mitochondrial carrier proteins is dependent on the GET pathway. Life Sci. Alliance. 4. doi:10.26508/lsa.202000918.

Yogev, O., and O. Pines. 2011. Dual targeting of mitochondrial proteins: mechanism, regulation and function. Biochim. Biophys. Acta. 1808:1012–1020. doi:10.1016/j.bbamem.2010.07.004.

